# Septin-Mediated RhoA Activation Engages the Exocyst Complex to Recruit the Cilium Transition Zone

**DOI:** 10.1101/2022.05.20.492811

**Authors:** Darya Safavian, Moshe S. Kim, Hong Xie, Lu Dai, Richard F. Collins, Carol Froese, Koh-ichi Nagata, William S. Trimble

## Abstract

Septins are filamentous GTPases that play important, but poorly characterized roles in ciliogenesis. Here, we show that SEPTIN9 regulates RhoA signaling at the base of cilia by binding and activating the RhoA guanine nucleotide exchange factor, ARHGEF18. GTP-RhoA is known to activate the membrane targeting exocyst complex, and suppression of SEPTIN9 causes disruption of ciliogenesis and mislocalization of an exocyst subunit, SEC8. Using basal body-targeted proteins we show that upregulating RhoA signaling at the cilium can rescue ciliary defects and mislocalization of SEC8 caused by global SEPTIN9 depletion. Moreover, we demonstrate that transition zone components, RPGRIP1L and TCTN2 fail to accumulate at the transition zone in cells lacking SEPTIN9 or depleted of the exocyst complex. Thus, SEPTIN9 regulates the recruitment of transition zone proteins on Golgi-derived vesicles by activating the exocyst via RhoA to allow the formation of primary cilia.

## INTRODUCTION

Primary cilia are non-motile protrusions of the plasma membrane found on nearly every vertebrate cell and they are formed from microtubules that emanate from membrane-docked basal bodies (Garcia-Gonzalo & Reiter, 2012). These organelles sense and transduce diverse signals to regulate cell physiology, development and organ homeostasis (Gerdes et al., 2007). Mutations disrupting ciliary structure and function cause a wide range of human disorders collectively known as ciliopathies that affect nearly all organs (Sharma et al., 2008). In order to relay extracellular cues into the cell, both the lipid and protein compositions of ciliary membranes are distinct from those of the contiguous cytosol and plasma membrane. This specialized composition of the cilium has led to suggestions that a diffusion barrier may exist at the base of cilia to prevent mixing of these membrane domains. Septins were initially suggested to form a ring-like membrane diffusion barrier near the base of cilia (Hu et al., 2010) although subsequent studies revealed that they were not always present in a ring at the base of cilia (Ghossoub et al., 2013). Septins comprise a novel component of the cytoskeleton and are filament-forming GTPases that, in addition to their role in ciliogenesis, have been implicated in diverse cellular processes including cytokinesis, cell migration and phagocytosis (Ghossoub et al., 2013; Hu et al., 2010; Huang et al., 2011). Subsequent studies have shown that depletion of SEPTIN2 resulted in the failure to assemble the transition zone (Chih et al., 2012) and this may explain the contribution of septins in ciliary membrane diffusion. However, the mechanism by which septins influence transition zone assembly is not known.

The transition zone functions as a ciliary gate during early stages of ciliogenesis to control the ciliary entry and exit of proteins to maintain the unique cilia composition. A B9D1-containing macromolecular complex comprised of more than 10 transition zone proteins, including Tectonic 1 (TCTN1), TCTN2, CC2D2A, and TMEM231 (Chih et al., 2012; Garcia-Gonzalo et al., 2011) are recruited early during the formation of the transition zone. Mutations in B9D1 genes have been revealed to cause ciliopathy-associated Meckel syndrome (MKS) (Hopp et al., 2011). Furthermore, a protein complex, consisting of three nephronophthisis (NPHP)-associated proteins, NPHP1, NPHP4 and RPGRIP1L localizes to the transition zone at the base of the cilium (Sang et al., 2011) and mutations in these components are linked to ciliopathies. The MKS and NPHP complexes are essential for normal cilia function, acting as a diffusion barrier to maintain the cilia membrane as a compartmentalized signaling organelle.

Septins have been linked to a large multi-subunit macromolecular tethering complex called the exocyst (Mei & Guo, 2018; Wu & Guo, 2015). The complex comprises eight subunits (Sec3, Sec5, Sec6, Sec8, Sec10, Sec15, Exo70, Exo84) and is mainly involved in the tethering of post-Golgi secretory vesicles to sites of exocytosis in the cell (Guo et al., 1999; Novick et al., 1980; TerBush et al., 1996). Sec3 and Exo70 bind to phosphoinositide lipids through their PH domains and are thought to provide a membrane patch or ‘landmark’ at the plasma membrane for the formation of the complex. In contrast, the remaining exocyst subunits arrive at the exocytosis site along with the secretory vesicle (He et al., 2007; Yamashita et al., 2010). Interestingly, both Sec3 and Exo70 bind to GTP-Rho in yeast, indicating that these exocyst components are effectors of RhoA (Guo et al., 2001; Wu et al., 2010). Although the precise interaction of septins with the exocyst complex are ill-defined, septins are known to recruit the exocyst complex to active membrane remodeling sites in the growing bud of yeast, during mammalian cytokinesis and in the formation of a penetrating peg in pathogenic rice fungi (Barral et al., 2000; Estey et al., 2010; Gupta et al., 2015). Like septins, the exocyst complex has been previously shown to be necessary for ciliogenesis and exocyst subunits have been localized to the ciliary base as well as the axoneme (Seixas et al., 2016; Zuo et al., 2009).

Although previous studies have implicated multimeric complexes such as septins, exocyst, MKS and NPHP in ciliogenesis (Chih et al., 2012; Ghossoub et al., 2013; Hu et al., 2010; Sang et al., 2011; Seixas et al., 2016; Zuo et al., 2009), the relationship between these components and the related signaling pathway driving ciliogenesis was not known. Here, we describe an unexpected regulatory role for septins in RhoA signaling and exocyst recruitment at the cilia basal body and have found that the exocyst complex functions to deliver transition zone proteins to the ciliary base for the formation of the transition zone and concomitant cilia formation.

## RESULTS

### SEPTIN9 binds ARHGEF18 and Promotes RhoA Activation

Previously, ARHGEF18 was identified as a SEPTIN9 interacting protein in a yeast two-hybrid screen (Nagata & Inagaki, 2005) and the interaction was mapped to the N-terminus of SEPTIN9. To confirm this interaction occurs in retinal pigment epithelium cells, hTERT-RPE1 cells were transfected with Flag-SEPTIN2, Flag-SEPTIN6, Flag-SEPTIN7, and Flag-SEPTIN9, individually (Supplementary Figure 1). hTERT-RPE1 cells were also co-transfected with Myc-ARHGEF18, Flag-RhoA, Flag-SEPTIN9, and the heterotypic septin complex (Supplementary Figure 1 and Figure 1). All constructs were expressed at approximately the same level individually in h TERT-RPE1 cells and the mobilities of each band were confirmed in singly transfected cells (Supplementary Figure 1). As we previously noted, we found that Myc-tagged ARHGEF18 bound to His6-SEPTIN9, but not to the SEPTIN9ΔN mutant (Supplementary Figure 2A). Next, we asked whether SEPTIN9 and ARHGEF18 could form a stable protein complex with RhoA. His6-tagged RhoA was able to form a ternary complex with ARHGEF18 and SEPTIN9 but could not bind to SEPTIN9 alone (Supplementary Figure 2B). Thus, the association of RhoA with SEPTIN9 was dependent on the presence of ARHGEF18. To determine if the same association might occur when SEPTIN9 was in a prototypical septin complex along with SEPTIN2, SEPTIN6, and SEPTIN7, we co-expressed SEPTIN2, SEPTIN6, SEPTIN7 and SEPTIN9 with RhoA and ARHGEF18. His6-RhoA was able to pull-down the septin complex and ARHGEF18 but could not bind the septin complex alone (Supplementary Figure 2C).

**Figure 1.**
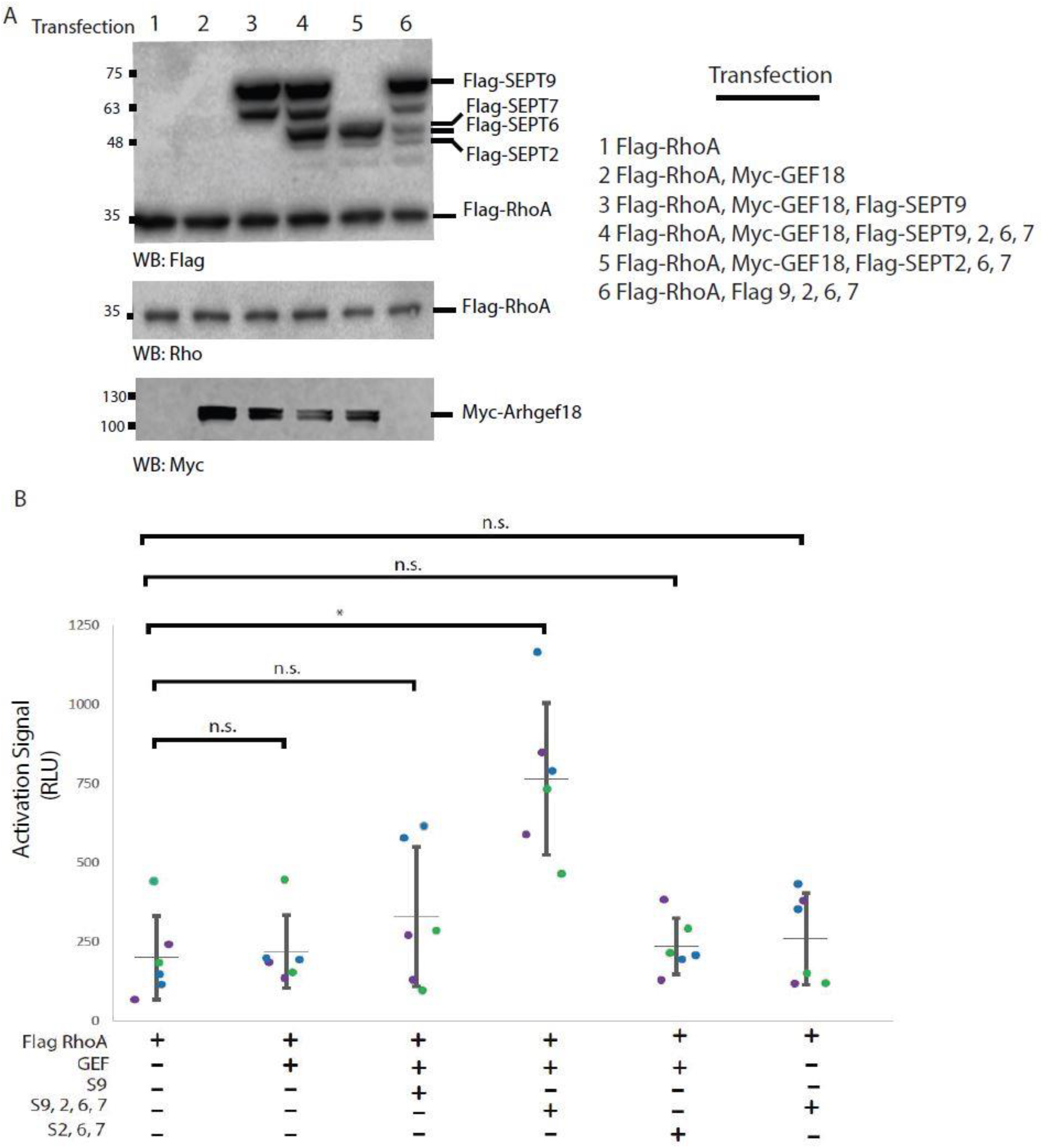
hTERT-RPE1 cells were co-transfected with Myc-ARHGEF18, Flag-His6-RhoA and/or a heterotypic 2-6-7-9 septin complex and septin9. Cell lysates were incubated with glutathione-agarose coupled to GST-RhoA binding domain (GST-RBD) to determine levels of GTP-bound Flag-His6-RhoA. All input lanes represent cell lysates and were immunoblotted with Flag and Rho antibodies. (B) Quantitation of active GTP-RhoA in cells transfected with ARHGEF18, RhoA and/or a heterotypic 2-6-7-9 septin complex and Septin9 using G-lisa RhoA Activation Assay Kit (Cytoskeleton Inc). The data show relative luminescence units (RLU) over background signal (wells of the Rho-GTP affinity plate incubated with lysis buffer alone instead of cell lysates). Cells transfected with all three components, including Myc-Arhgef18, Flag-His6-RhoA and a heterotypic 2-6-7-9 septin complex had the largest increase in GTP-bound RhoA. This represents 3 independent experiments, and asterisk denotes Student’s t-test p-value significance *=p<0.05. n.s. No significance is denoted.

Given the stable interactions between ARHGEF18, RhoA and the septin complex, we asked if the septin complex might regulate the activity of RhoA. To do so, the levels of Rho activity were measured using RhoA G-lisa activation assay, wherein the Rho-GTP-binding domain (RBD) of a Rho effector is coupled to a microwell plate, allowing affinity-based detection of the active Rho in samples. Compared to RhoA alone, expression of ARHGEF18 in hTERT-RPE cells resulted in a small increase in the levels of active RhoA (Figure 1B). Surprisingly, when the heterotypic septin complex (SEPTIN2, SEPTIN6, SEPTIN7 and SEPTIN9) was transfected with ARHGEF18 and RhoA, there was a significantly larger increase in the levels of active RhoA, compared to SEPTIN9 alone (Figure 1B). To rule out the possibility that the septin complex might be the only component responsible for this level of RhoA activation, lysates expressing the septin complex alone did not appreciably increase the levels of GTP-RhoA (Figure 1B). Thus, expression of both ARHGEF18 and the septin complex together had a synergistic effect on RhoA activation that could not be achieved by either ARHGEF18 or the septin complex alone.

Previous studies had suggested that SEPTIN9 binding inhibited ARHGEF18 activity (Nagata & Inagaki, 2005), but in those studies SEPTIN9 alone was overexpressed, so we speculate that the inhibitory effect in that case may have resulted from expression of SEPTIN9 that is not stoichiometrically associated with septin complexes or filaments. Some DH-PH-containing proteins are known to regulate their own activity through intramolecular inhibition (Bi et al., 2001; Yu et al., 2010). This autoinhibition is relieved by proteins binding to the inhibitory domain of the guanine nucleotide exchange factor. The C-terminus of ARHGEF18 has been predicted to act as its autoinhibitory domain (Tsuji et al., 2010) and this region has also been shown to interact with the N-terminal variable region of SEPTIN9 (Nagata & Inagaki, 2005). We hypothesize that the effect of septin complexes on ARHGEF18 is mediated by direct interaction of SEPTIN9 and ARHGEF18, which relieves the autoinhibition on its nucleotide exchange domain. Expression of SEPTIN9 as part of the complex or filament may increase activation by increasing the avidity of the interaction with the ARHGEF18. The DH-PH domain is the active domain of RhoGEF, and in isolation is constitutively active (Schmidt & Hall, 2002). Indeed, expression of the N terminal DH-PH region alone was sufficient to activate RhoA. At similar protein expression levels compared to full length ARHGEF18, the DH-PH domain induced an abundant array of actin stress fibers in HeLa cells, similar to a constitutively active RhoA mutant (Supplementary Figure 3). Septin-mediated Rho activation is also observed in yeast. A fission yeast RhoGEF, GEF3, has been shown to interact with the septin complex and activate Rho4 during cytokinesis (Wang et al., 2014). It would be of interest to determine whether septin-mediated activation of RhoGEFs in fission yeast and mammalian cells occur by a conserved mechanism involving relief of RhoGEF autoinhibition.

### SEPTIN9 and ARHGEF18 localize to the Base of Primary Cilia

Active RhoA localizes to the base of cilia in multiciliated cells and its localization is dependent on components of the planar cell polarity (PCP) pathway (Pan et al., 2007; Park et al., 2008).

Dishevelled is a core component of the PCP pathway that also regulates ciliogenesis by recruiting GTP-RhoA to the base of Xenopus cilia (Park et al., 2008). ARHGEF18 was previously found to interact with Dishevelled in a neuronal screen for RhoA activation (Tsuji et al., 2010). Since both GTP-RhoA and Dishevelled are localized to the base of cilia, we asked if ARHGEF18 might localize to the base of cilia in a cell line commonly used for primary cilia study, hTERT-RPE1. Indeed, immunostaining with antibody specific for ARHGEF18 showed that it localized as two puncta at the base of primary cilia (Figure 2A and Supplementary Figure 4A). These puncta localized to basal bodies as revealed by their overlap with gamma-tubulin (Figure 2B).

**Figure 2.**
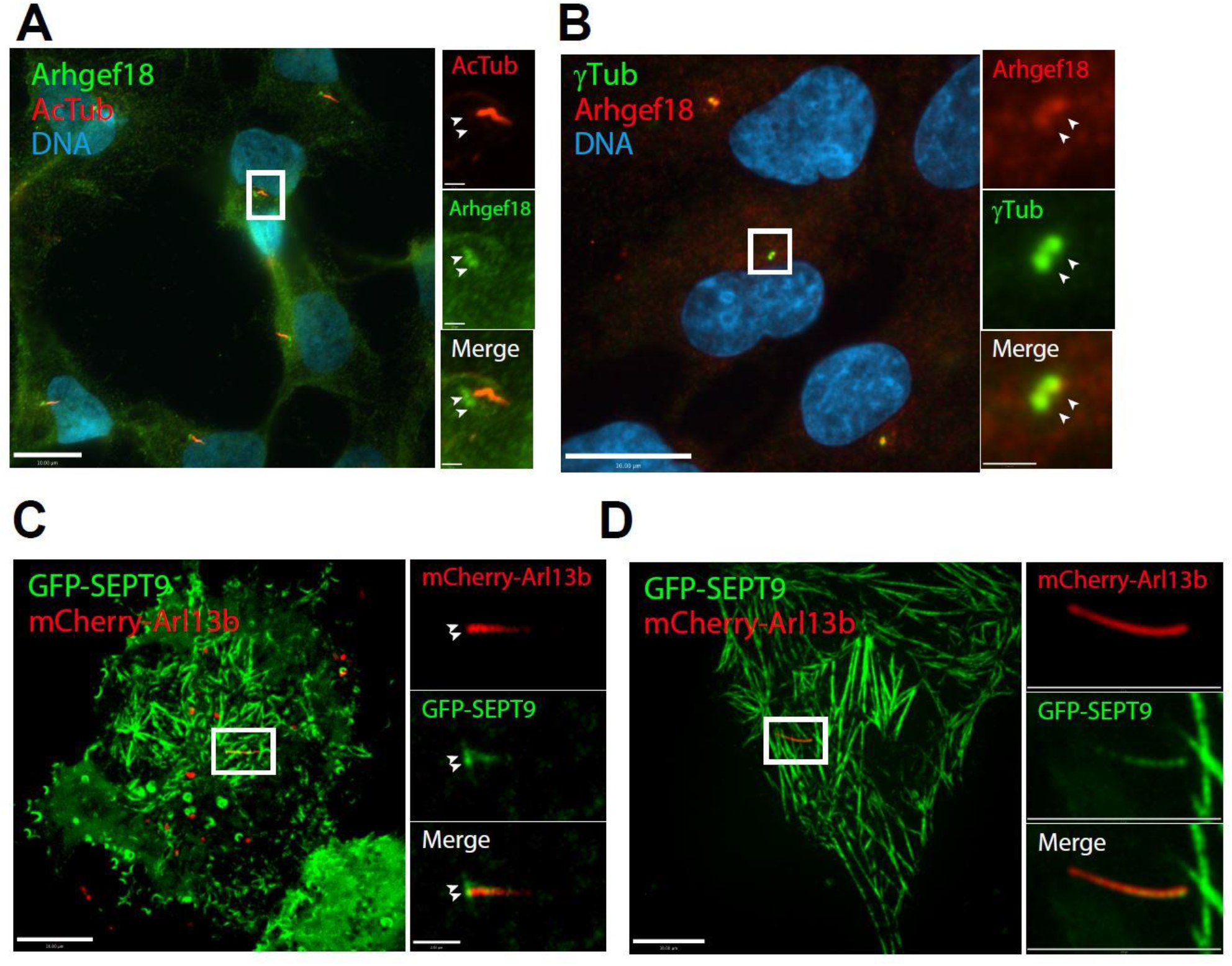
SEPTIN9 and ARHGEF18 localize to the base of primary cilia (A and B) ARHGEF18 localizes to the base of primary cilia in hTERT-RPE1 cells. (A) ARHGEF18 localizes as two puncta at the base of cilia (arrowhead). Cells were PFA-fixed and cilia were stained with acetylated tubulin antibody (AcTub, red). DNA was counterstained with Hoescht 33342 (blue). Bar represents 10 μm. Boxed region is magnified on right. Bar in magnified images represent 1μm. (B) ARHGEF18 associates with gamma-tubulin (γTub, arrowhead) in serum-starved hTERT-RPE1 cells fixed in ice-cold methanol. Bar represents 10 μm. Boxed region is magnified on right. Bar in magnified images represent 1μm. (C and D) Live-cell microscopy of hTERT-RPE1 co-transfected with GFP-SEPTIN9-i3 (green) and mCherry-Arl13b (red, an axonemal marker) at 16h serum starvation. (C) GFP-SEPTIN9-i3 localises as two foci at the base of cilia and the lower part of the axoneme (right panel). Boxed regions are magnified. Magnified regions show only the ciliary Z-planes, whereas the unmagnified images show all Zplanes. Bar represents 10 μm. Boxed region is magnified on right. Bar in magnified images represent 2μm. (D) Live-cell microscopy of hTERT-RPE1 co-transfected with GFP-SEPTIN9-i3 and mCherry-Arl13b at 24h serum starvation. GFP-SEPTIN9-i3 co-localises to the lower two thirds of the axoneme. Bar represents 10 μm. Boxed region is magnified on right. Bar in magnified images represent 2μm.

Next, we examined if septins might also localize to the base of cilia. Previous studies have shown the localization of mammalian septins at either the base of cilia or along the length of cilia, so we tried to determine conditions where septins might localize to the base of cilia. Consistent with another report (Ghossoub et al., 2013), we found that septins were predominantly distributed along the length of cilia by immunostaining of fixed RPE cells (Figure 2C, and unpublished data). However, when we examined ciliary localization of GFP-tagged septins under live-cell conditions, we detected GFP-SEPTIN9 at two distinct locations: along the length of cilia and/or at the base of cilia (Figure 2C and 2D). Furthermore, septin localization at the base of cilia appeared as two puncta consistent with the basal body and daughter centriole (arrowheads in Figure 2C). Taken together, septins localize to both the basal bodies and along the length of cilia raising the possibility that septins might have multiple functions within cilia.

### SEPTIN9 or ARHGEF18-depleted Cells Show Ciliary Defects

To determine if SEPTIN9 and ARHGEF18 are required for the formation of cilia, we individually depleted either SEPTIN9 or ARHGEF18 (Figures 3A and 3E). Compared to controls, SEPTIN9 depletion did not affect ciliation frequency, but resulted in a decrease in ciliary length (Figure 3B-D). This is consistent with previously reported ciliary defects in SEPTIN9-depleted cells (Ghossoub et al., 2013). The defects associated with ARHGEF18 depletion were more drastic, showing a loss of two-thirds of cilia and those few cilia that remained were short (Figure 3F-H). The differences in the severity of cilia defects might arise from basal activity of ARHGEF18 in the SEPTIN9-depleted cells that would not be present in ARHGEF18-depleted cells or could be due to residual SEPTIN9 following depletion. We therefore generated a SEPTIN9 knockout cell line using CRISPR/Cas9 to eliminate SEPTIN9 and better address its role in ciliogenesis. Our immunofluorescence studies revealed that the absence of SEPTIN9 leads to severe cilia loss (Figure 3J-L). The difference in the severity of cilia defect is likely due to the partial depletion of SEPTIN9 in siRNA-treated cells versus the complete absence in CRISPR cell line (Figure 3I). Nonetheless, it is evident that SEPTIN9 and ARHGEF18 are required for the formation of cilia and/or the regulation of ciliogenesis.

**Figure 3.**
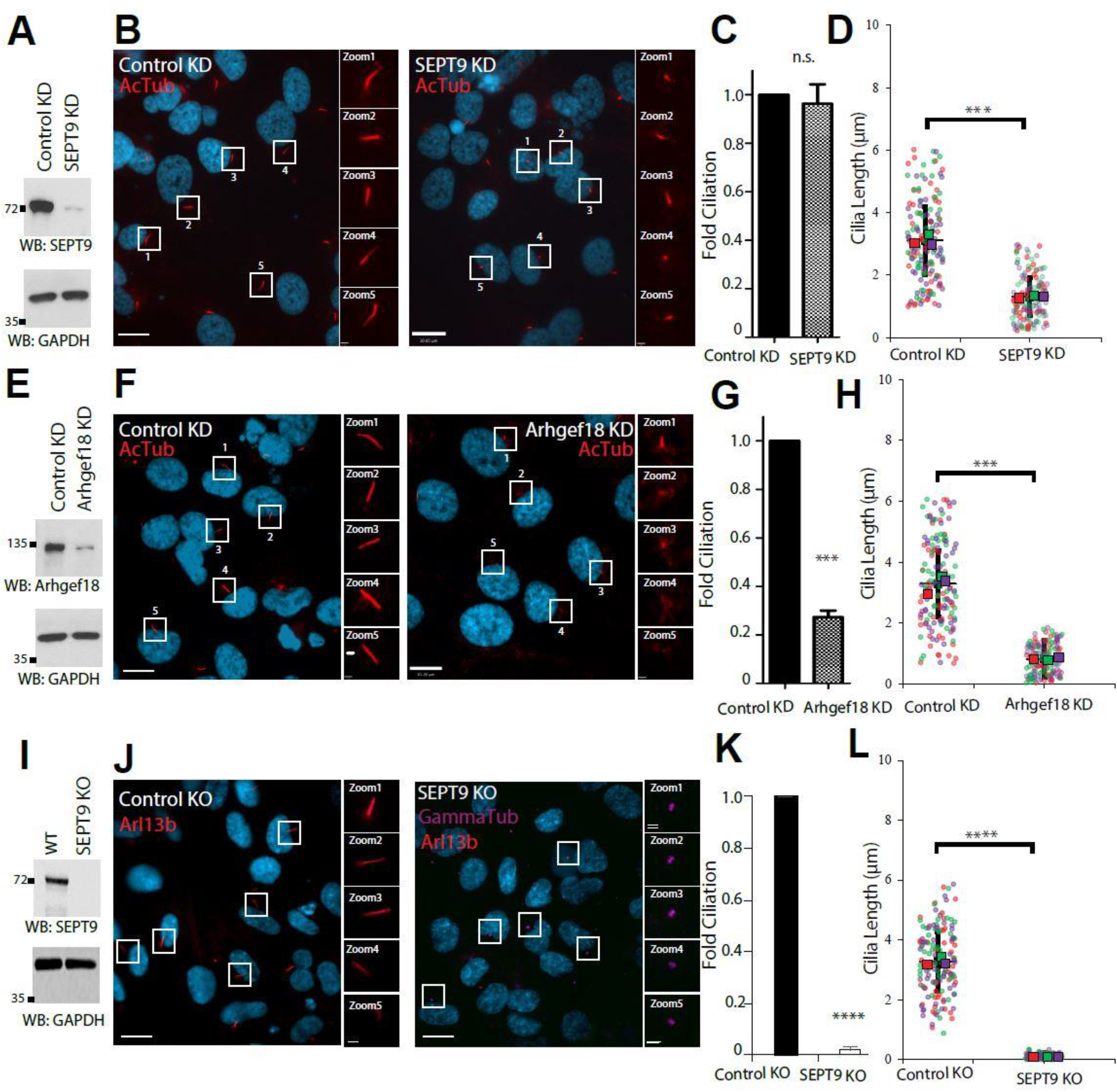
SEPTIN9 or ARHGEF18-depleted cells show ciliary defects. (A-D) SEPTIN9 regulates ciliary length in serum-starved hTERT-RPE1 cells. (A) Western blot showing depletion of SEPTIN9 protein with siRNA treatment. (B) Cilia are shorter in SEPTIN9-depleted cells. Cilia are immunostained with acetylated tubulin antibody (AcTub). Bar represents 10 μm. Bars in magnified images represent 1 μm. (C and D) Quantification of ciliation and ciliary length in SEPTIN9-depleted cells. Cells depleted of SEPTIN9 still form cilia, but they are shorter. Three independent experiments were conducted and 100 cells were counted in each experiment. (E-H) ARHGEF18 is required for the formation of cilia. (E) Western blot showing depletion of ARHGEF18 protein with siRNA treatment. (F) Cilia are absent or severely stunted in ARHGEF18-depleted cells. (G and H) Quantification of ciliogenesis defect in ARHGEF18-depleted cells. No significance is denoted n.s. (I-L) SEPTIN9 is required for the formation of cilia. (I) Western blot showing deletion of SEPTIN9 with CRISPR/Cas9. (J) Cilia are absent in SEPTIN9 deleted cells and the few cilia that are formed are severely stunted. (K and L) Quantification of ciliogenesis defect in SEPTIN9-knockout cells. Data are represented as mean ± SEM; three independent experiments were conducted and 100 cells were counted in each experiment. Student’s t-test p-value significance *=p<0.05, **=p<0.01, ***=p<0.001 and ****=p<0.0001. n.s. No significance is denoted.

### Basal body-associated ARHGEF18 and SEPTIN9 Regulate Ciliary Length

In serum-starved hTERT-RPE1 cells, SEPTIN9 localizes to diverse cytoskeleton structures including actin stress fibers and distinct ciliary compartments. With the establishment of the biological function of SEPTIN9 in ciliogenesis, we then sought to understand the relationship between SEPTIN9 and RhoA signaling at the base of cilia. To determine if SEPTIN9 at the basal body is involved in the regulation of ciliogenesis, we attempted to rescue ciliogenesis defects in SEPTIN9 knockout cells by targeting SEPTIN9 to the basal body with a GFP-Centrin-SEPTIN9 fusion protein. Centrin is a centrosome-associated protein that is localized in the basal body at the base of the cilium and immunostaining with an anti-GFP antibody demonstrated correct targeting of centrin and GFP-tagged versions of it, to the base of cilia. This basal body-targeted SEPTIN9 construct was observed to rescue the ciliogenesis defects in SEPTIN9 knockout cells (Figure 4E and 4F). Since ARHGEF18 binds the N-terminus of SEPTIN9 (Nagata & Inagaki, 2005), we tested if the fusion of the N-terminal 300 amino acids, containing the 147 amino acids of SEPTIN9 shown to bind ARHGEF18 to GFP-Centrin is sufficient to rescue ciliogenesis.

**Figure 4.**
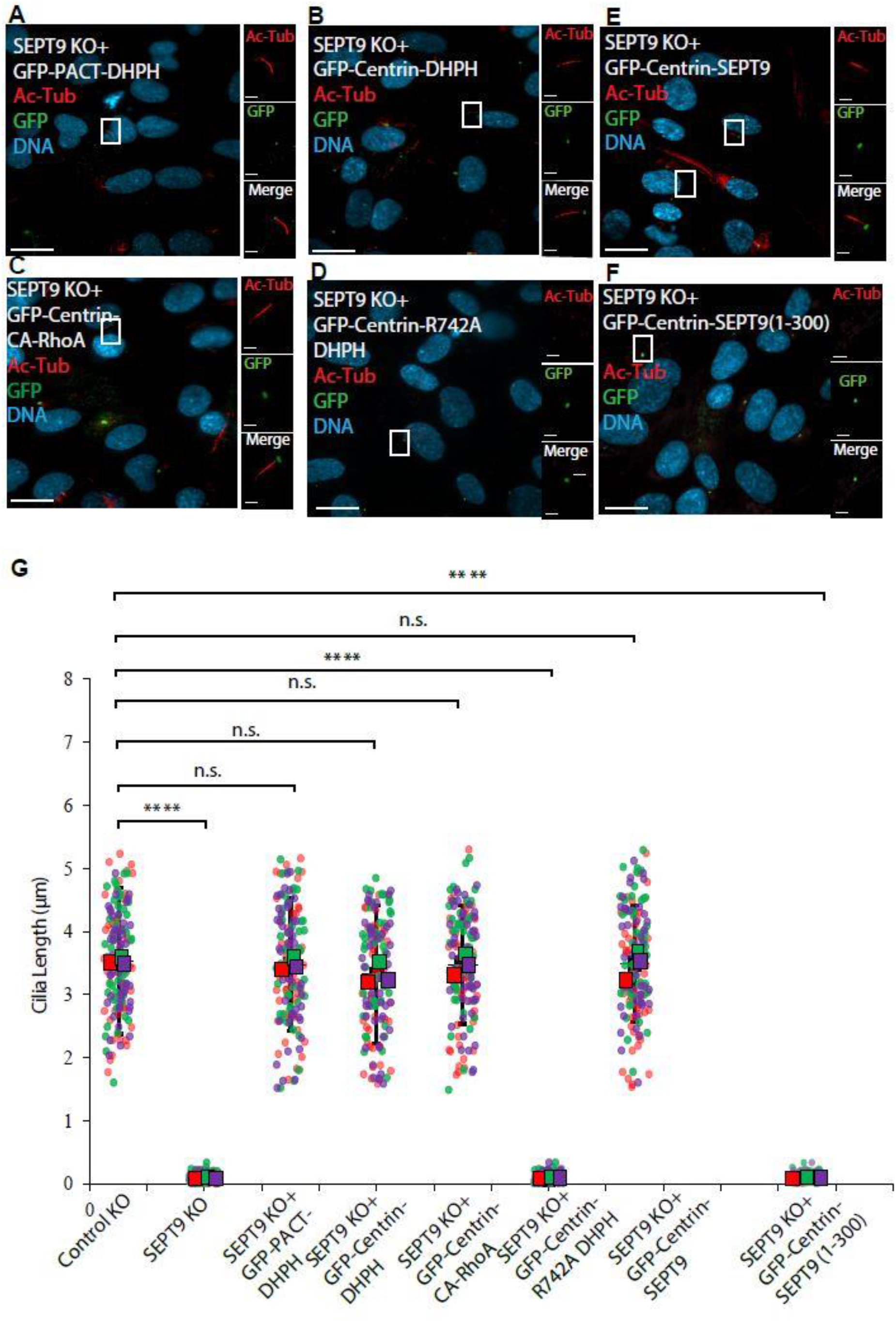
The basal body targeting of DH-PH domain rescued ciliogenesis in SEPTIN9 knockout cells. (A, B, C, E) The PACT as well as Centrin-driven basal body targeting of the DH-PH domain and the CA-RhoA restored ciliogenesis in SEPTIN9 knockout cells. (D) The DH-PH mutation that blocks the interaction with RhoA failed to rescue ciliogenesis. (E) The basal body expression of SEPTIN9 restored cilia formation in knockout cells. (F) The basal body expression of the N-terminal fragment of SEPTIN9 failed to rescue cilia formation. (G) Quantification of ciliary length in Control knockout cells, SEPTIN9 knockout cells, SEPTIN9 knockout cells expressing GFP-PACT-DH-PH, GFP-Centrin-DH-PH, mutant GFP-Centrin-R742A DH-PH, GFP-Centrin-CA-RhoA, GFP-Centrin-SEPTIN9, and GFP-Centrin-SEPTIN9 (1-300), respectively. Data are represented as mean ± SEM; the data achieved from 3 independent experiments for each cell line and 100 cells were counted in each experiment. Student’s t-test p-value significance *=p<0.05, **=p<0.01, ***=p<0.001 and ****=p<0.0001. n.s. No significance is denoted. Boxed regions are magnified. Bar represents 10 μm. Bars in magnified images represent 1 μm.

However, it was found that the basal body expression of the N-terminal region of SEPTIN9 alone could not rescue ciliogenesis defects caused by the loss of SEPTIN9 in knockout cells (Figure 4F). This observation is consistent with our G-lisa data (Figure 1) where, compared to SEPTIN9 alone, the expression of SEPTIN9, as a component of the heterotypic complex, along with SEPTIN 2, 6, 7, and GEF18 had a synergistic effect on RhoA activity. Taken together, these data strongly suggest that SEPTIN9 as part of a septin complex at the basal body is involved in the regulation of ciliogenesis and provide the first evidence for a RhoA mediated signaling pathway promoted by septins.

The interface between the DH-PH domain of RhoGEF and RhoA has been characterized, and point mutation of a specific argenine at the binding site between the DH-PH domain of RhoGEF and RhoA to alanine greatly reduces the nucleotide exchange activity (Chen et al., 2010). To test the significance of the GEF in RhoA activation on ciliogenesis, a R742A mutation was generated in an N-terminal GFP and Centrin tagged (GFP-Centrin-R742A DH-PH) construct. As expected, the mutant DH-PH expressed at the basal body failed to rescue the ciliary length and ciliogenesis defects in SEPTIN9 KO cells (Figure 4D and 4F).

To confirm that the activation of RhoA was not related to the centrin targeting tag, we fused GFP-PACT to the DH-PH domain of ARHGEF18. The PACT domain (pericentrin-AKAP-450 controsomal targeting) represents a coiled coil region close to the C-terminus of centrosomal proteins that serves to recruit AKAP-450 and pericentrin to the controsome. Thus, fusion of the PACT domain to another protein confers centrosomal targeting. In SEPTIN9 knockout cells, the PACT-mediated basal body targeting of DH-PH also rescued ciliogenesis and restored ciliary length (Figure 4A and 4F). Similarly, the expression of GFP-Centrin-DH-PH and GFP-Centrin-Constitutively Active (CA) RhoA in SEPTIN9 knockout cells rescued ciliogenesis and restored the ciliary length (Figure 4B, 4C, 4F). Strikingly, the global expression of DH-PH in SEPTIN9 KO cells did not rescue ciliogenesis, indicating that the targeted expression of DH-PH at the ciliary base is essential to restore the formation of primary cilia.

Next, we asked whether SEPTIN9 and ARHGEF18 might act in the same pathway to regulate ciliary length. As septins act as molecular scaffolds, we tested whether SEPTIN9 was required for the recruitment of ARHGEF18 to basal bodies. The loss or depletion of SEPTIN9 did not result in the mislocalization of ARHGEF18 away from the basal bodies (Supplementary Figure 4B). In contrast, ARHGEF18 was found to be required for the axonemal localisation of SEPTIN9 (Supplementary Figure 4A). This suggests that the axonemal accumulation of septins depends on their binding to ARHGEF18 located at the basal body. We believe that this interaction and basal body co-localization is transient and is not easily detected in fixed cells while the axonemal localization of septins is stable. We further investigated the axonemal localization of other septins upon the expression of GFP-Centrin-DH-PH in SEPTIN9 Knock out cells and found that the localization of SEPTIN7 and SEPTIN2 in the rescued ciliary axoneme also depends on the basal body association of SEPTIN9 and ARHGEF18 since loss of SEPTIN9 also eliminates axonemal septin accumulation in the cilia (Supplementary Figure 5).

Finally, we tested if impairing or potentiating RhoA signaling at the base of cilia in wild type cells might regulate cilia length and to test this GFP-Centrin was fused to dominant negative (DN) RhoA. Compared to untransfected cells (Figure 5A and 5E), or cells expressing only GFP-Centrin (Figure 5B and 5E), the cilia were considerably shorter in cells expressing basal body targeted DN RhoA (Figure 5C and 5E). Moreover, in wild type ciliated cells, the centin-derived basal body targeting of the constitutively active DH-PH led to an increase in the ciliary length (Figure 5D and 5E). Thus, tuning of activity of RhoA signaling at the base of cilia could increase or decrease ciliary length compared to controls.

**Figure 5.**
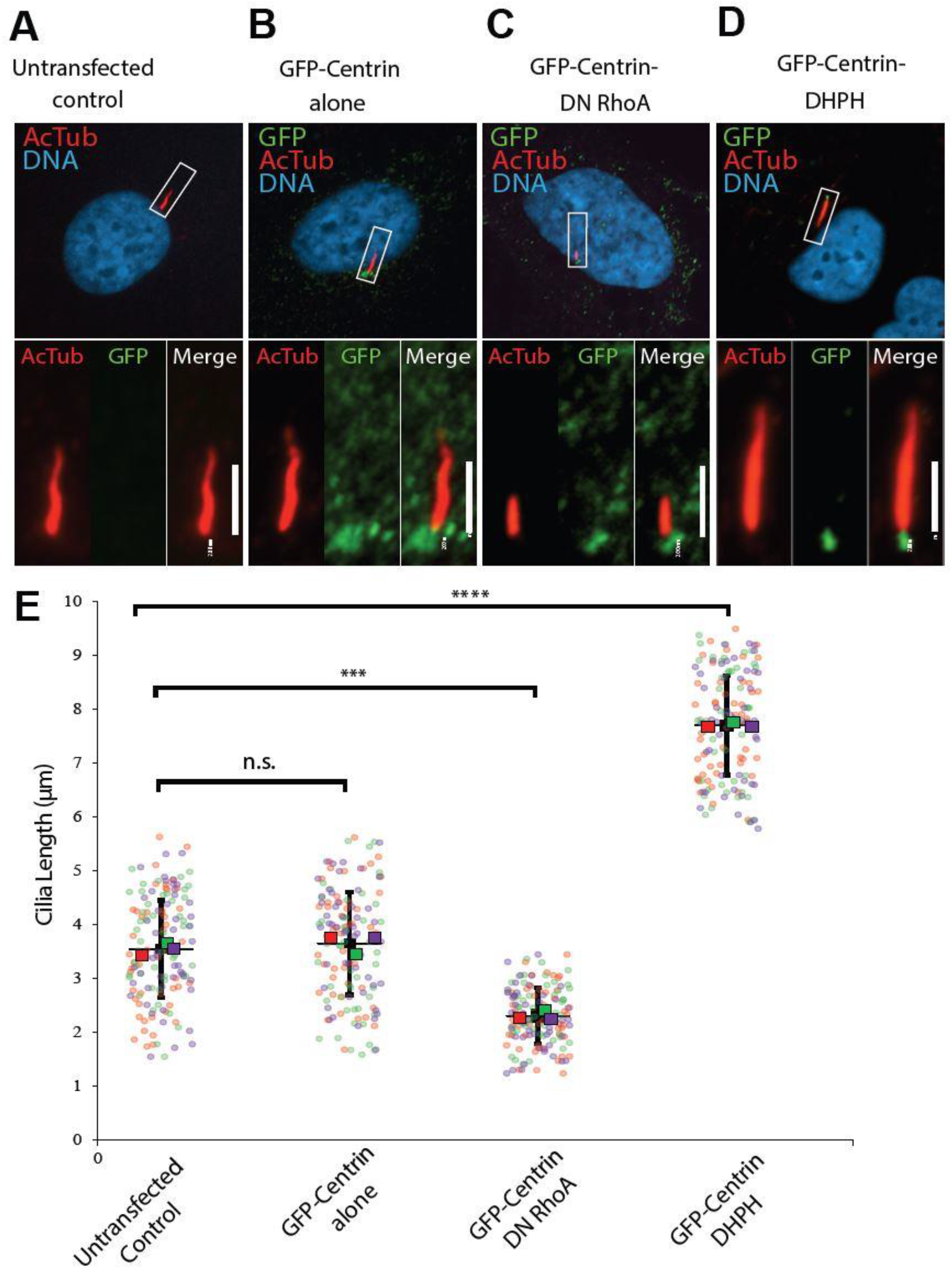
Basal body-associated SEPTIN9 and ARHGEF18 regulate ciliary length. Serum-starved hTERT-RPE1 cells were fixed in methanol and stained with GFP and acetylated tubulin (AcTub) antibodies. Targeted RhoA signaling at the base of cilia and effects on ciliary length. (C) DN-RhoA was targeted to the base of cilia, resulting in shorter cilia compared to either vector alone (B) or untransfected cells (A). (D) The DH-PH domain of ARHGEF18 was targeted to the base of cilia, resulting in longer cilia compared to controls (A and B). (E) Quantification of ciliary length in WT hTERT-RPE1 cells expressing GFP alone, GFP-Centrin-DN RhoA, and GFP-Centrin-DH-PH. Data are represented as mean ± SEM; three independent experiments were conducted and 100 cells were counted in each experiment. Student’s t-test p-value significance *=p<0.05, **=p<0.01, ***=p<0.001 and ****=p<0.0001. n.s. No significance is denoted. Boxed regions are magnified. Magnified bar represents 2μm.

### Basal Body recruitment of Exocyst component SEC8, but not EXO70 Depends on SEPTIN9

Having established that SEPTIN9 and ARHGEF18 are both essential for proper cilia formation and that the SEPTIN9-ARHGEF18 interaction is required to activate RhoA, we set out to identify effectors downstream of RhoA with the intention of further defining the molecular mechanisms driving ciliogenesis. RhoA is known to control the localization and assembly of the exocyst complex, which closely associates with septins (Guo et al., 2001; Hsu et al., 1998; Wu et al., 2010). In addition, GEF-H1, a RhoA GEF highly related to ARHGEF18, was shown to facilitate exocyst assembly and secretion in other systems (Pathak et al., 2012). We therefore examined the function of SEPTIN9 in recruiting the exocyst complex to the ciliary base. Previous studies showed exocyst components localizing at basal bodies (Seixas et al., 2016; Zuo et al., 2009). Using confocal microscopy, we analyzed the localization of endogenous exocyst subunits in hTERT-RPE1 cells. Consistent with previous studies in other systems, both the lardmark component EXO70 and the vesicle associated component SEC8 were localized at the base of the mature cilia in wild type cells (Figures 6A and 6E). Similarly, EXO70 was also detected colocalizing with gamma tubulin at the basal body in SEPTIN9 knockout cells (Figure 6A). By contrast, deletion of SEPTIN9 resulted in a drastic loss of SEC8 signal from the basal body (Figure 6F). EXO70 functions as a spatial landmark at the plasma membrane for the assembly of the exocyst complex, which in turn tethers secretory vesicles for exocytosis (Boyd et al., 2004). Thus, the conserved basal body localization of EXO70 in mature cilia in WT and SEPTIN9 knockout cells suggests that the localization of the landmark EXO70, along with SEC3, is independent of SEPTIN9. In contrast to EXO70, SEC8, along with the other five subunits, is transported to the sites of the exocyst assembly via secretory vesicles (Boyd et al., 2004; Finger & Novick, 1997). Therefore, the impaired localization of the SEC8 subunit to the basal body in SEPTIN9 knockout cells would be consistent with SEPTIN9 functioning to promote localized exocyst assembly and vesicle docking, a RhoA-dependent process (Pathak et al., 2012). To further confirm the role of SEPTIN9 in activating RhoA and subsequent exocyst assembly at the base of the cilia, we tested the possibility of rescuing exocyst recruitment by expressing a GFP-Centrin-DH-PH construct to locally activate RhoA. As predicted, the expression of GFP-Centrin-DH-PH in the SEPTIN9 knockout (KO) cells rescued the defect and restored both ciliogenesis and SEC8 localization to the base of cilia (Figures 6G and 6H). Thus, SEPTIN9, through interaction with ARHGEF18, activates RhoA at the basal body to facilitate exocyst assembly concomitant with vesicle tethering and cargo delivery to the site of ciliogenesis.

**Figure 6.**
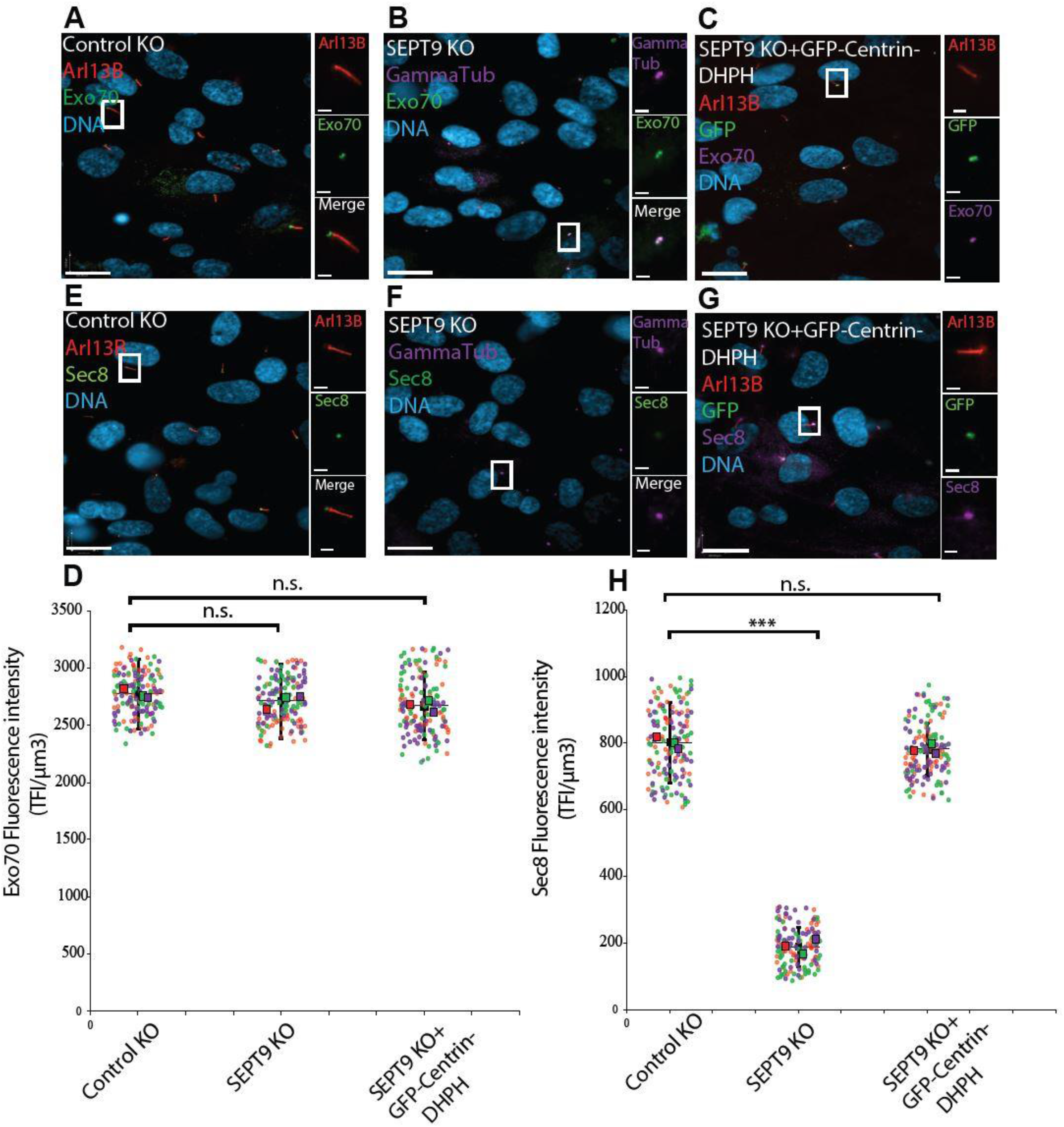
SEPTIN9 and Activated RhoA recruit exocyst components to the basal body and regulate ciliogenesis. Confluent serum-starved hTERT-RPE1 cells were fixed in methanol and stained with anti-SEC8, EXO70, ARL13B, gamma tubulin and GFP antibodies. (A and E) Exocyst subunits are localized at the base of cilia in hTERT-RPE1 cells. (A) In wild-type (WT) hTERT-RPE1 cells, EXO70, the spatial landmark of the exocyst on the target membrane (green), and (E) SEC8, the vesicle associated subunit (green), are localized to the base of cilia marked by ARL13B (red). (B and F) In SEPTIN9 CRISPR knockout cells, EXO70 (green) localized to the ciliary base stained by gamma tubulin (purple) (B), while the absence of SEPTIN9 resulted in a drastic loss of SEC8 (green) at the basal body (purple) (F). (C and G) Rescue of SEPTIN9 knockout cells using a basal body-targeting construct expressing the DH-PH of ARHGEF18. The expression of DH-PH domain of ARHGEF18 at the base of cilia via the GFP-Centrin-DH-PH construct in SEPTIN9 knockout cells rescued SEC8 (green) localization and restored the formation of cilia stained by ARL13B (red) (G). Similar to wild type (A) and SEPTIN9 knockout cells (B), EXO70 localized to the ciliary base in the rescue experiment (C). (D and H) Quantification of Exo70 and Sec8 signal intensity in Control knockout cells, SEPTIN9 knockout cells, and SEPTIN9 knockout cells expressing GFP-Centrin-DH-PH. In each image, all pixel values were in the range of 0-255 and no under or over saturated pixels were detected. A Region of Interest (ROI) of 98.74 µm^3^ (containing 24 pixels) was selected at the base of the cilia, marked by Gamma Tubulin, in all the quantifications of the fluorescence intensity of Exo70 and Sec8 proteins. The same ROI was used to measure the background fluorescence, subtracted from the Sum of ROI fluorescence reads for each protein (TFI/µm^3^; Telomere Fluorescence Intensity/µm^3^). Data are represented as mean ± SEM, achieved from three independent experiments and 100 cells were counted in each experiment. Student’s t-test p-value significance *=p<0.05, **=p<0.01, ***=p<0.001 and ****=p<0.0001. n.s. No significance is denoted. Bar represents 10 μm. Bars in magnified images represented 1 μm.

### The Localization of Transition Zone Components is Dependent on SEPTIN9 and the Exocyst Complex

With the establishment of the biological function of the septins in recruiting the exocyst to the ciliary base, we then investigated which ciliary cargo might be targeted to primary cilia in an exocyst-and septin-dependent manner. Previously, multisubunit complexes have been identified at the transition zone and shown to be crucial for both the biogenesis and function of the cilia

(Chih et al., 2012; Garcia-Gonzalo et al., 2011; Huang et al., 2011; Sang et al., 2011). Loss of function of these transition zone complexes, containing Merkel Syndrome (MKS), nephronophthisis (NPHP) and CEP290 proteins have been associated with compromised ciliogenesis in some tissues, and deregulated ciliary protein composition in others, leading to numerous clinical symptoms as well as early lethality (Czarnecki & Shah, 2012). However, how the transition zone components are delivered to the base of the primary cilium was largely unknown. Since septins were previously shown to be involved in the localization of transition zone proteins Tmem231, B9D1 and Cc2d2a (Chih et al., 2012), we focused our efforts on representative components of the two distinct complexes that localize to the transition zone.

Using antibodies specific for ciliopathy proteins RPGRIP1L, a member of the NPHP complex and Tectonic2 (TCTN2), a component of the B9D1 complex, we find that both are localized to the transition zone in a SEPTIN9-dependent manner. In control cells, we consistently observed that TCTN2 was localized to a single position between the axonemal microtubules (MTs) and basal bodies, when co-immunostained for acetylated tubulin and gamma tubulin, respectively (Figure 7E). RPGRIP1L localized to three distinct puncta at the base of the cilium. Two of these puncta colocalized with gamma tubulin, consistent with RPGRIP1L being a centrosomal protein, while the third punctum localized at the transition zone (Figure 7A). RPGRIP1L is known to be required for the transition zone localization of all MKS module proteins in a cell type-specific manner, indicating it occupies a core position within the assembly (Shi et al., 2017; Wiegering et al., 2018; Williams et al., 2011; Yang et al., 2015). Deletion of SEPTIN9 resulted in loss of the transition zone-localized RPGRIP1L signal with a clear shift from 3 to 2 puncta, while the TCTN2 localization is absent in SEPTIN9 knockout cells (Figure 7B, 7K, 7F and 7L). The mislocalization of these transition zone proteins was rescued by the basal body-specific targeting of DH-PH, using a centrin targeting construct encoding GFP-Centrin-DH-PH (Figures 7C, 7K, 7G, 7L). Similar to SEPTIN9 knockout cells, SEC8-depleted cells had short cilia and impaired localization of the transition zone proteins at the base of the cilium (Figures 7D, 7L, 7H).

**Figure 7.**
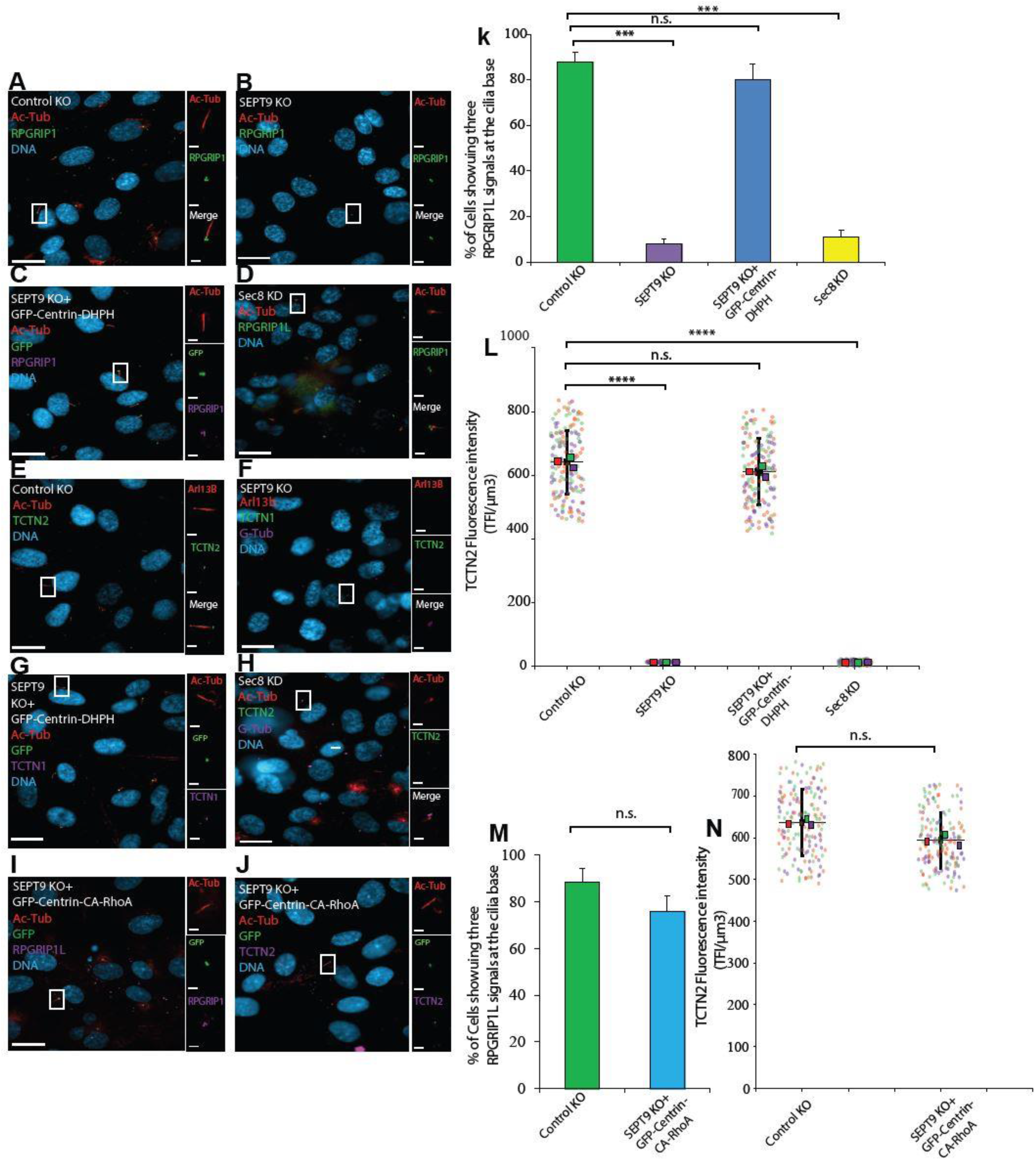
SEPTIN9, Exocyst, and ARHGEF18 mediate the localization of transition zone components to the ciliary base. Confluent serum-starved hTERT-RPE1 cells were fixed in methanol and stained with anti-RPGRIP1L, TCTN1, ARL13B, acetylated tubulin (Ac-Tub), and GFP. (A and B) WT and SEPTIN9 knockout hTERT-RPE1 cells were stained with representatives of two distinct complexes at the transition zone. (A) RPGRIP1L (green) appeared as three distinct puncta, localized at the base of the cilium stained with Ac-Tub (red). (B) In SEPTIN9 CRISPR knockout cells, the absence of SEPTIN9 resulted in impaired localization of transition zone components, with a clear shift in the RPGRIP1L signal from three to two puncta. (D) Like SEPTIN9 suppression, the depletion of Sec8 in SiRNA knockdown cells, resulted in the mislocalization of the RPGRIP1L signal from the transition zone. (C) The basal body expression of DH-PH domain of ARHGEF18 in SEPTIN9 knockout cells via the GFP-Centrin-DH-PH construct rescued the localization of RPGRIP1L at the base of cilia stained with ARL13B (red).. (E and F) In WT hTERT-RPE1 cells, TCTN2 is localized at the transition zone, while the absence of SEPTIN9 results in a drastic mislocalization of TCTN2 in Knockout cells. (G) The basal body expression of DH-PH domain of ARHGEF18 in SEPTIN9 knockout cells via the GFP-Centrin-DH-PH construct rescued the localization of TCTN2 at the base of cilia stained with ARL13B (red). (H) Similar to SEPTIN9 knockout cells, the depletion of SEC8 subunit of the exocyst complex impaired the localization of TCTN2 (green) away from the basal body stained with gamma-tubulin (purple). (I and J) The basal body expression of constitutively active RhoA in SEPTIN9 knockout cells restored the RPGRIP1L and TCTN2 localization, as well as ciliogenesis. (k) Percentage of cells showing three RPGRIP1L signals at the ciliary base in the Control knockout cells, SEPTIN9 knockout cells, SEPTIN9 knockout cells expressing GFP-Centrin-DH-PH, and SEC8 siRNA depleted cells. (L) Quantification of TCTN2 fluorescence intensity in Control knockout cells, SEPTIN9 knockout cells, SEPTIN9 knockout cells expressing GFP-Centrin-DH-PH, and SEC8 siRNA depleted cells. (M) Percentage of cells showing three RPGRIP1L signals at the ciliary base in Control knockout cells, and SEPTIN9 knockout cells expressing GFP-Centrin-CA-RhoA. (N) Quantification of TCTN2 fluorescence intensity in Control knockout cells, and SEPTIN9 knockout cells expressing GFP-Centrin-CA-RhoA. In each image, all pixel values were in the range of 0-255 and no under or over saturated pixels were detected. A Region of Interest (ROI) of 98.74 µm^3^ (containing 24 pixels) was selected at the base of the cilia in all the quantifications of fluorescence intensity of TCTN2. The same ROI was used to measure the background fluorescence, subtracted from the Sum of ROI fluorescence reads for TCTN2 (TFI/µm^3^; Telomere Fluorescence Intensity/µm^3^). Data are represented as mean ± SEM; three independent experiments were conducted and in each experiment 100 cells were counted. Student’s t-test p-value significance *=p<0.05, **=p<0.01, ***=p<0.001 and ****=p<0.0001. n.s.=No significance is denoted. Bar represents 10 μm. Bars in magnified images represents 1 μm.

Remarkably, the expression of the constitutively active RhoA at the base of cilia in SEPTIN9 knockout cells restored the localization of transition zone proteins and rescued ciliogenesis (Figures 7I, 7J, 7M and 7N). Taken together, this data supports the conclusion that SEPTIN9, through activating RhoA, mediates the localization of the vesicle-tethering exocyst complex to the base of the cilium, promoting the delivery of transition zone components to form the ciliary gate.

**Figure 8.**
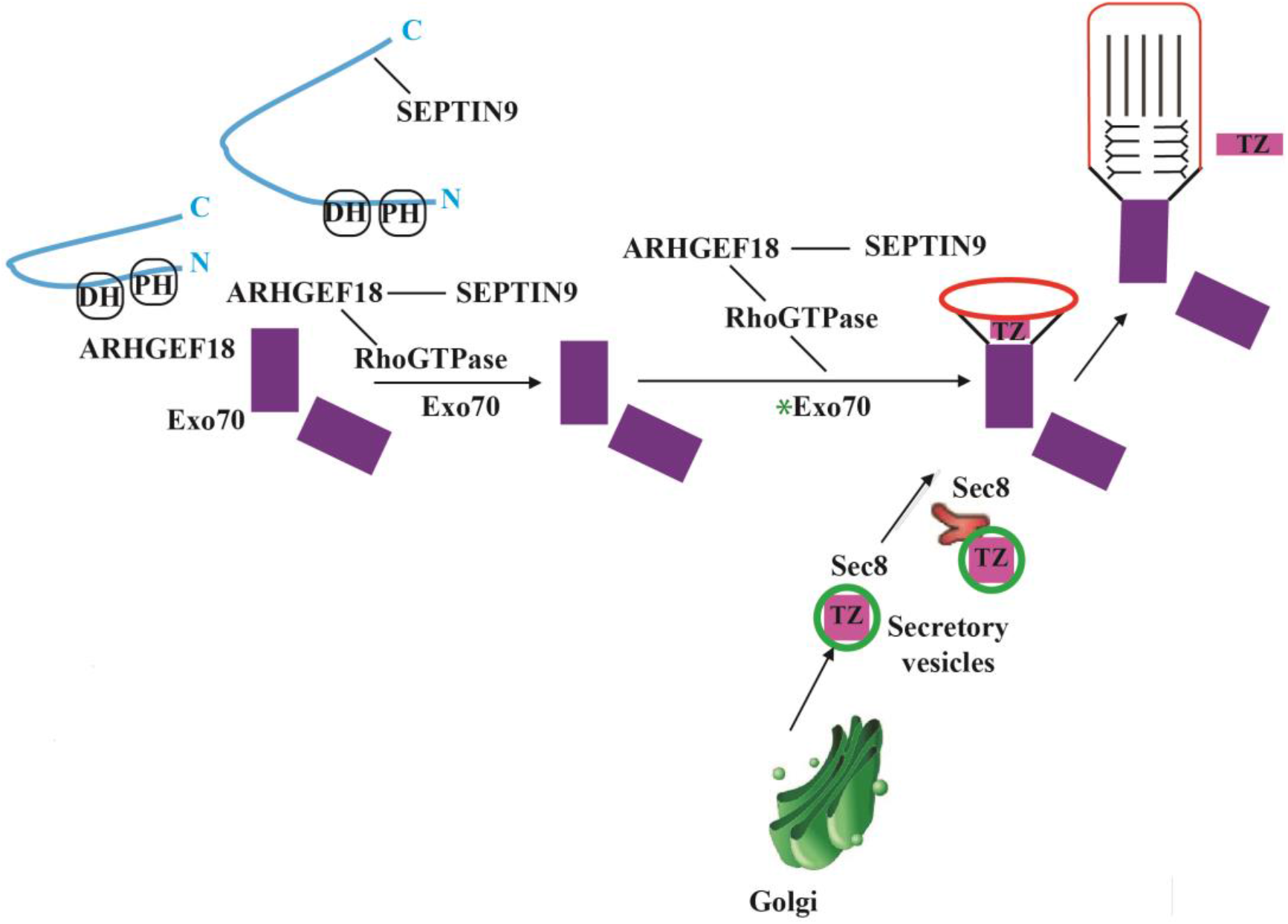
Proposed Model of SEPTIN9-mediated signaling pathway during early stages of ciliogenesis in hTERT-RPE1 cells. ARHGEF18 is localized at the basal body and is required for the localization of SEPTIN9 at the base of the primary cilium. The interaction between the N-terminus of SEPTIN9 and the C-terminus of ARHGEF18 is proposed to relieve the intramolecular autoinhibition of ARHGEF18 and results in the activation of the DH-PH domain. Subsequently, the activated DH-PH binds to and activates RhoAGTPase. Binding of RhoAGTPase to the Exo70 subunit (* indicates activated Exo70) of the exocyst activates this complex by blocking inhibitory interactions between Exo70 and other components of the complex, thus reorganizing the complex into a more active form. The activated exocyst complex then mediates the trafficking of post-Golgi derived secretory vesicles, tethered by the remaining six subunits including Sec8. These vesicles are suggested to deliver transition zone proteins (TZ) to the base of the cilia to promote ciliogenesis.

## DISCUSSION

Our data indicate that increasing or decreasing RhoA signaling at basal bodies can regulate ciliary length, and that a potential effector for RhoA in this context is the exocyst complex. The exocyst has previously been shown to be partitioned into two parts (Wu & Guo, 2015): landmark exocyst subunits (SEC3 and EXO70) bind to PIP2 on the plasma membrane which are thought to recruit vesicle-associated exocyst subunits (SEC5, SEC6, SEC8, SEC10, SEC15, EXO84) to the site of exocytosis. SEC3 and EXO70 are both known to bind to GTP-RhoA. The GTP-RhoA binding subsequently activates these landmarks at the basal body to facilitate the assembly of the entire exocyst complex, resulting in the delivery of post-Golgi vesicles and the subsequent cargo delivery (Adamo et al., 1999; Guo et al., 2001; Robinson et al., 1999; Zhang et al., 2001).

Previous studies demonstrated SEPTIN2-dependent recruitment of proteins (Cc2d2a, B9D1, Tmem231) to the transition zone, but the mechanism remained unclear (Chih et al., 2012). SEPTIN2 is a component of the core septin complex and its depletion likely affected SEPTIN9 function as well. Our data suggests a sequence of signaling events that ultimately lead to the correct targeting of transition zone proteins, RPGRIP1L and TCTN2. First, SEPTIN9 as part of a septin complex potentiates ARHGEF18 to activate RhoA, and RhoA-GTP then recruits the exocyst complex (Figure. 5), which in turn delivers vesicular cargo for the formation of the transition zone (Figure. 6).

At the ultrastructural level, the transition zone is comprised of Y-shaped macromolecular protein complexes that are thought to physically prevent the mixing of plasma membrane and ciliary membranes (Garcia-Gonzalo & Reiter, 2012). The tips of the Y-shape are predicted to be transmembrane or membrane-associated proteins, whereas the stem consists of proteins that are predicted to interact with the microtubule-rich axoneme. The use of fluorescence microscopy combined with electron microscopy revealed the localization of the TZ proteins including RPGRIP1L, TCTN2, and TMEM67 in human hTERT-RPE1 cells (Yang et al., 2015).

RPGRIP1L has been shown to occupy the most proximal position close to the basal body and transition fibers where it performs a central role as a scaffold for anchoring other MKS and NPHP module proteins in a cell type specific manner (Wiegering et al., 2018; Williams et al., 2011; Yang et al., 2015). The MKS complex component, TCTN2, is an extracellular protein, and presents at the same axial level as RPGRIP1L, but occupies the most peripheral position within the TZ structure (Yang et al., 2015). Additionally, functional interactions between different MKS and NPHP module proteins are found to be essential for anchoring the TZ components to the membrane during early stages of ciliogenesis (Williams et al., 2011). Since neither the cytoplasmic RPGRIP1L nor the extracellular TCTN2 were present, it seems likely that the entire TZ complex has failed to correctly assemble.

To date, it is not known precisely how or where these Y-shaped macromolecular protein complexes assemble. Several possibilities include an orderly assembly from the arms to the stem (outside-in), the stem to the arms (inside-out) or perhaps the entire Y-shaped complex comes pre-assembled. Both Golgi-derived and endosome-derived vesicles have been implicated in this process. Since the exocyst regulates post-Golgi vesicle traffic, this suggests that the Golgi-derived vesicles are required for the recruitment of TZ components arriving on endosomal vesicles. Further studies will be required to fully elucidate the role of the SEPTIN9-exocyst axis in transition zone protein recruitment and assembly.

## Acknowledgements

This work was supported by funds from CIHR fellowship MEF-158165 to DS and CIHR grant PJT-152194 to WST.

---

**Supplementary Figure 1.**
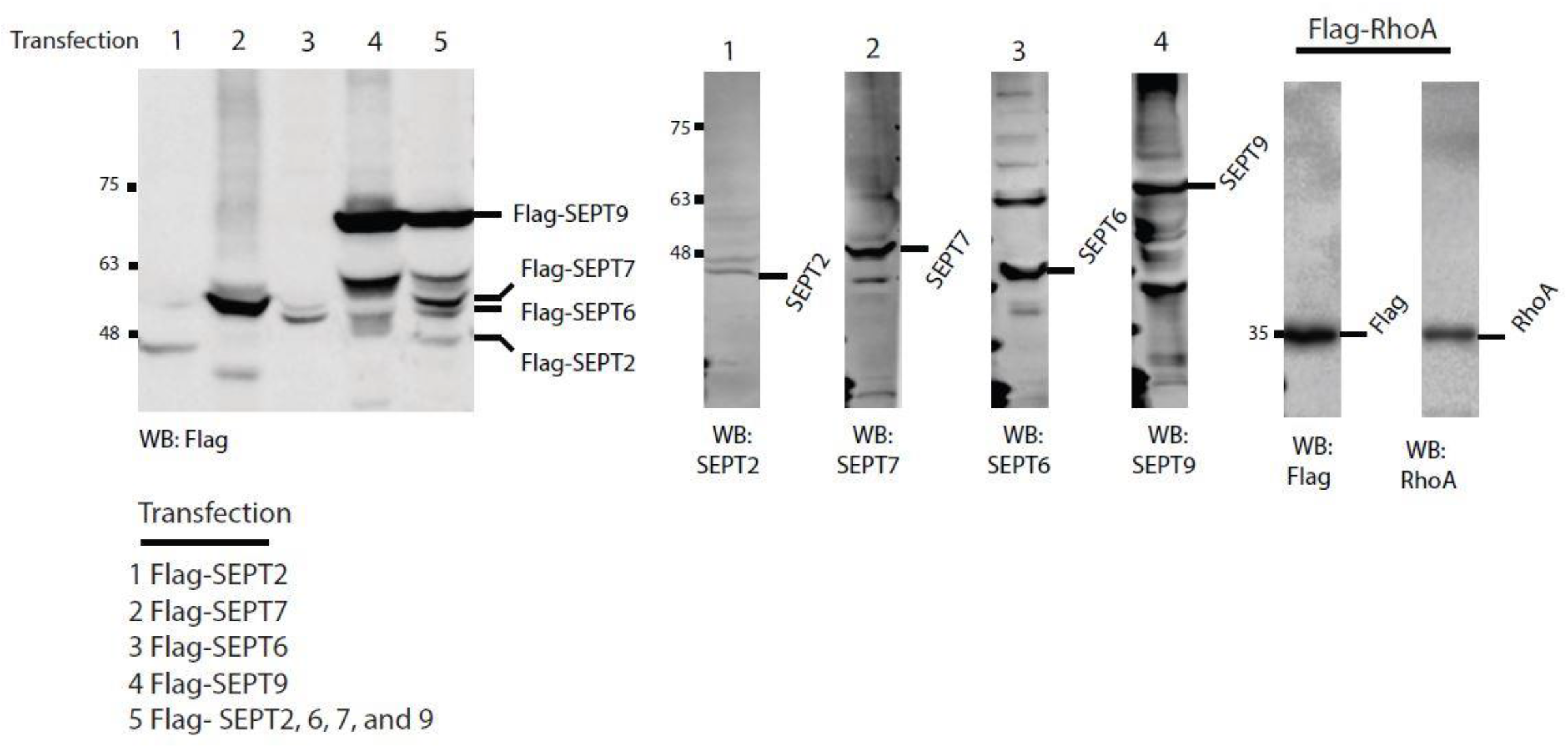
hTERT-RPE1 cells were transfected with Flag-SEPT2, Flag-SEPT6, Flag-SEPT7, Flag-SEPT9, a heterotypic 2-6-7-9 septin complex, and Flag-RhoA, respectively. All input lanes represent cell lysates and were immunoblotted with Flag, SEPT2, SEPT6, SEPT7, SEPT9, and Rho antibodies. All constructs were expressed at the same level individually in hTERT-RPE1 cells.

**Supplementary Figure 2.**
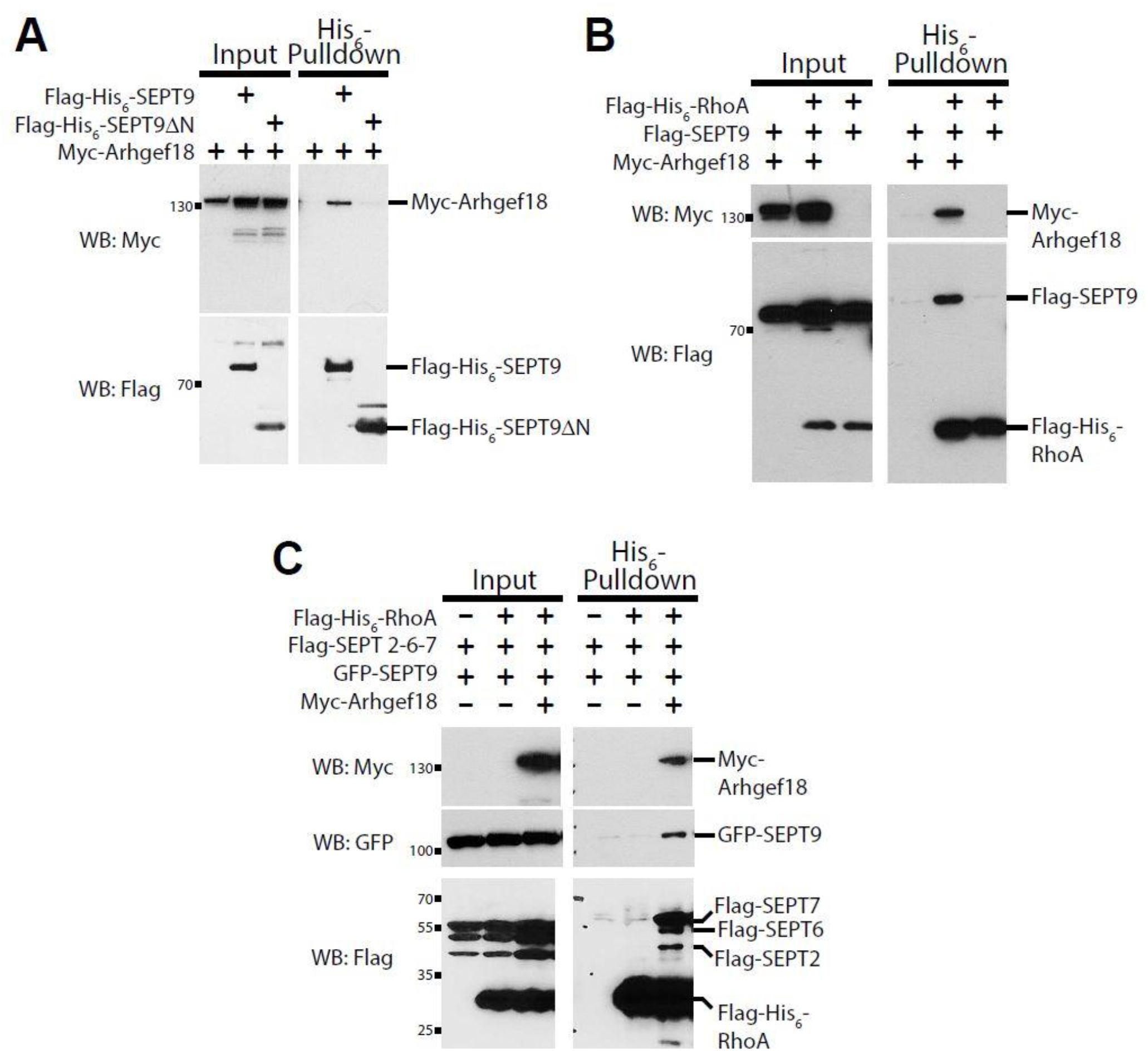
SEPTIN9 N-terminus binds ARHGEF18 (A) SEPTIN9 binds to ARHGEF18 through its N-terminus. HeLa cells were co-transfected with Myc-ARHGEF18 and either Flag-His6-SEPTIN9 or Flag-His6-SEPTIN9ΔN. Myc-Arhgef18 bound to SEPTIN9, but not to SEPTIN9ΔN. (B) SEPTIN9 forms a complex with RhoA and Arhgef18. HeLa cells were co-transfected with Myc-ARHGEF18, Flag-His6-RhoA and/or Flag-SEPTIN9. His6-RhoA was pulled down using nickel-agarose beads. SEPTIN9 bound to RhoA only in the presence of ARHGEF18. (C) A septin complex binds to Arhgef18 and RhoA. HeLa cells were co-transfected with Myc-ARHGEF18, Flag-His6-RhoA and/or a prototypical septin complex (comprising Flag-SEPTIN2, Flag-SEPTIN6, Flag-SEPTIN7 and GFP-SEPTIN9). The heterotypic septin complex bound to RhoA only in the presence of ARHGEF18.

**Supplementary Figure 3.**
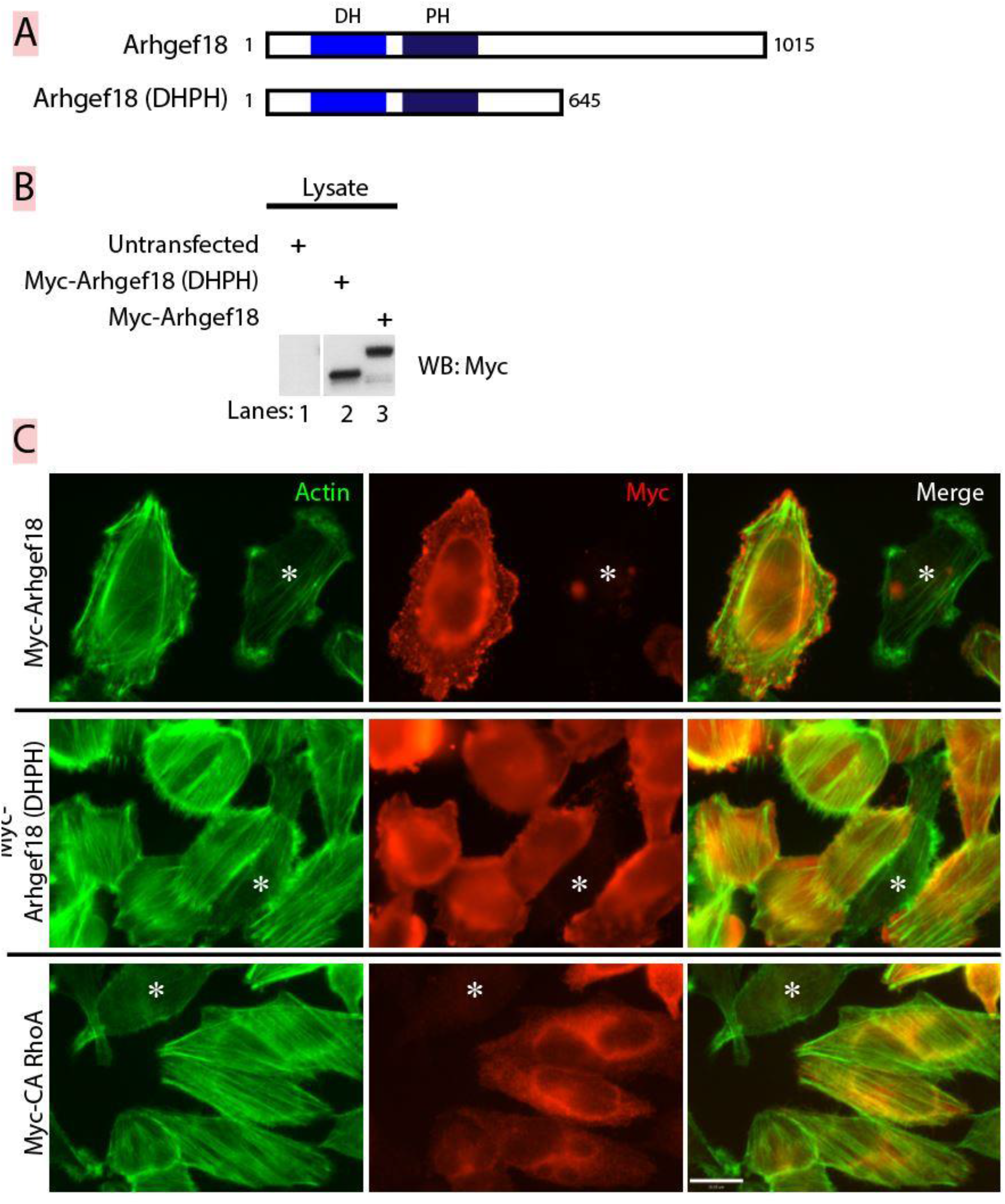
The DH-PH domain of ARHGEF18 stimulates formation of actin stress fibers (A) Domain structure of ARHGEF18. The DH-PH domain in relation to the full-length and a C-terminal truncation. (B) Expression of both full-length and DH-PH domain in HeLa cells. Both myc-tagged constructs are expressed in relatively equal amounts. (C) The DH-PH domain stimulates actin stress fiber assembly. Compared to untransfected cells (marked with an asterisk), full-length ARHGEF18 has a modest effect on increasing actin stress fibers. However, the DH-PH domain of ARHGEF18 shows a more pronounced increase in stress fiber assembly (compared to both untransfected cell [marked with an asterisk] and full-length ARHGEF18). The level of stress fiber induction is similar to expression of myc-tagged constitutively active RhoA (bottom panel). Bar represents 5 µm.

**Supplementary Figure 4.**
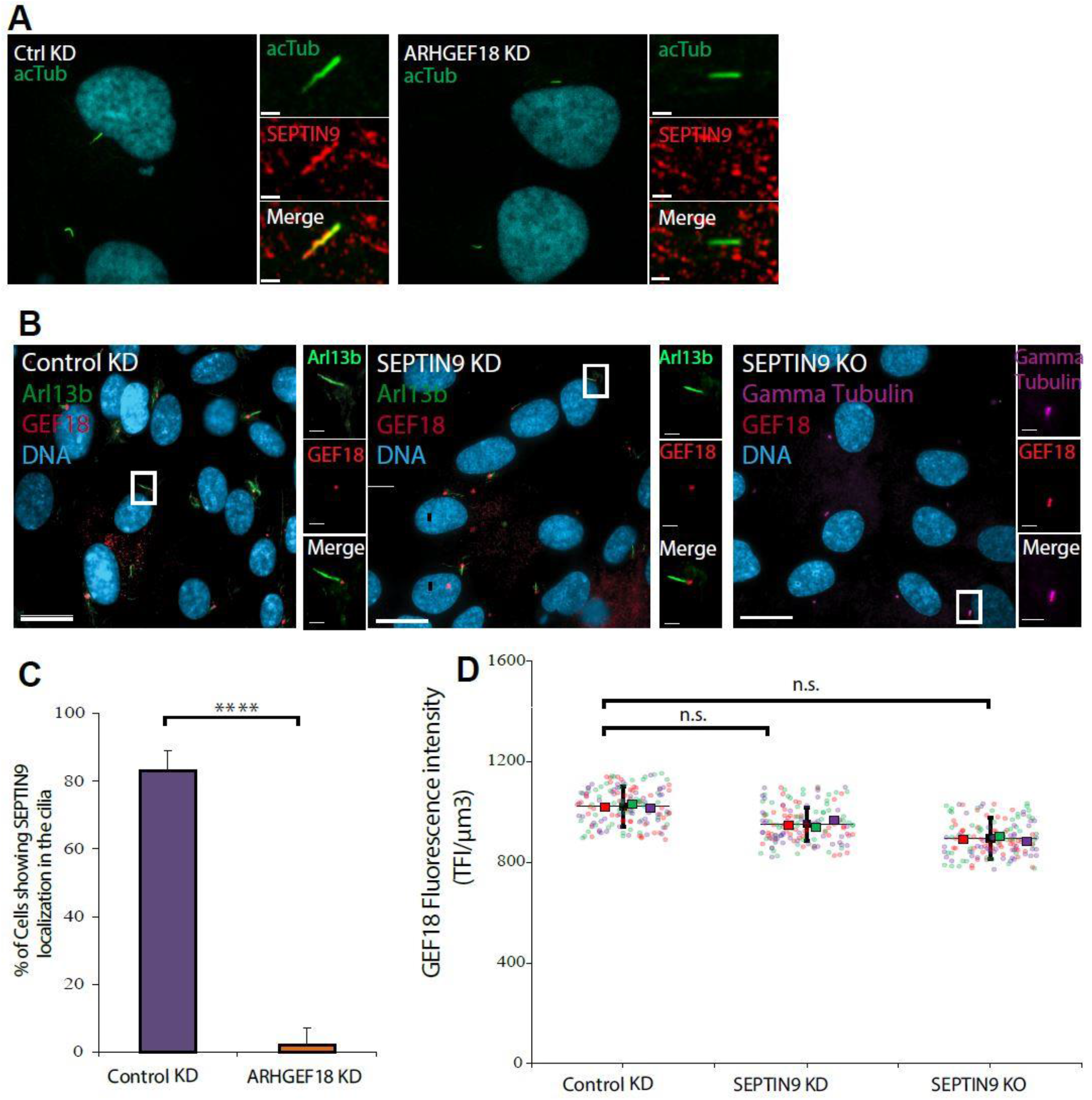
ARHGEF18 is required for ciliary localisation of SEPTIN9. (A) ARHGEF18 is required for ciliary localisation of SEPTIN9. Serum-starved hTERT-RPE1 cells depleted of ARHGEF18 were fixed and immunostained with SEPTIN9 (red) and acTub (green) antibodies. Ciliary localisation of SEPTIN9 is mislocalised in the rare cilia that still form when ARHGEF18 is depleted. (B) SEPTIN9 is not required for basal body localisation of ArhGEF18. Serum-starved hTERT-RPE1 cells depleted of SEPTIN9 and SEPTIN9 KO cells were fixed and immunostained with ARHGEF18 (red) and acTub (green, axonemal marker) antibodies. ARHGEF18 localisation at the base of cilia is unaffected with depletion or absence of SEPTIN9. (C) Quantification of percentage of cells showing SEPTIN9 localization in the axoneme in Control KD and ARHGEF18 KD cells. (D) Quantification of ARHGEF18 fluorescence intensity at the base of the cilia in Control KD, SEPTIN9 KD, and SEPTIN9 KO cells. Data are represented as mean ± SEM; three independent experiments were conducted and in each experimen100 cells were counted. Student’s t-test p-value significance *=p<0.05, **=p<0.01, ***=p<0.001 and ****=p<0.0001. n.s. No significance is denoted. Bar represents 10 μm. Bars in magnified images represents 1 μm.

**Supplementary Figure 5.**
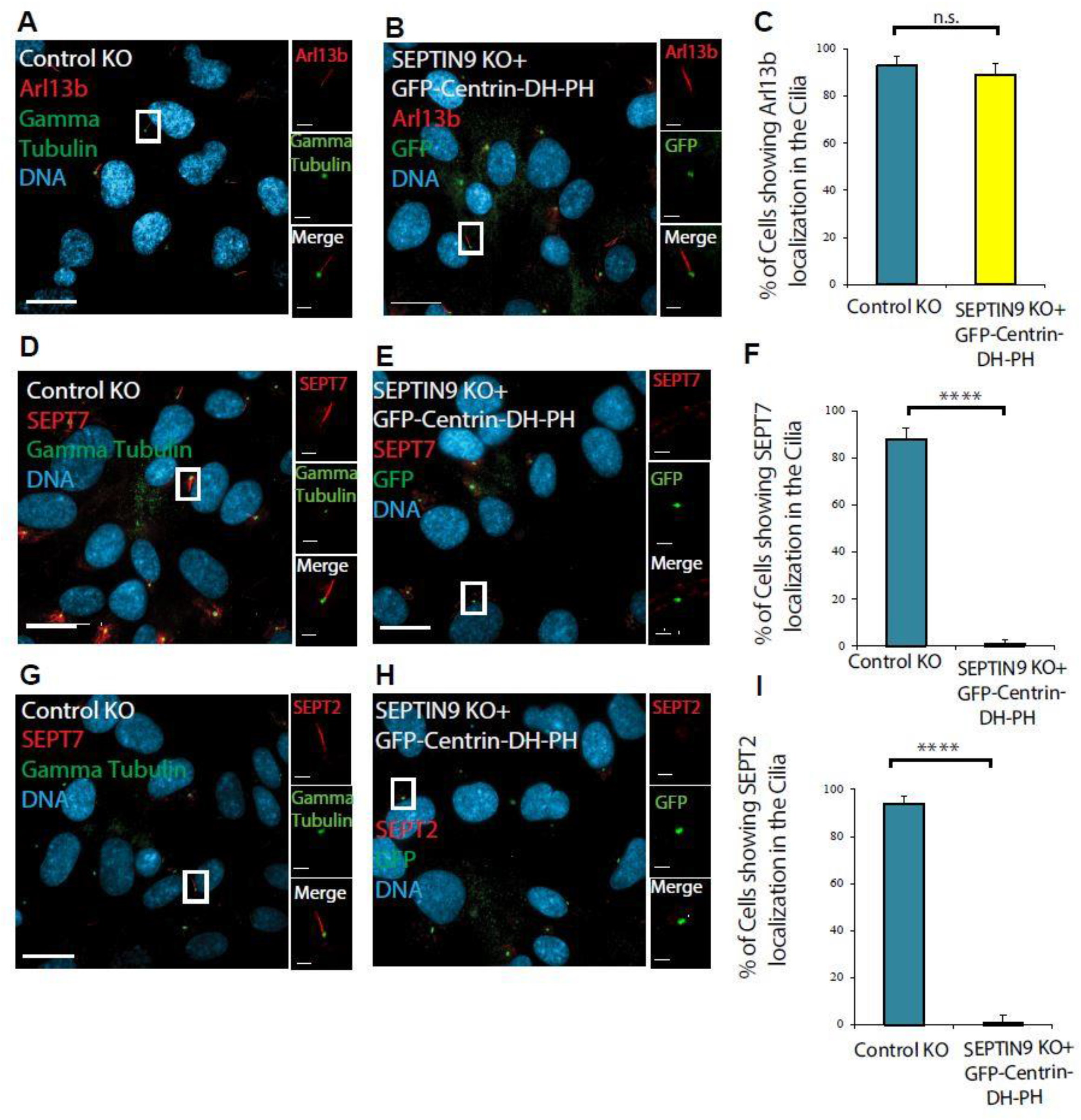
The axonemal septin accumulation depends on the SEPTIN9 and ARHGEF18 interaction and their transient basal body co-licalization. Serum-starved hTERT-RPE1 cells were fixed with PFA to visualize axonemal septin accumulation and stained with SEPTIN7, SEPTIN2, Gamma Tubulin, and GFP antibodies. (A and B) The expression of GFP-Centrin-DH-PH in SEPTIN9 KO cells rescued ciliogenesis in these cells measured by the percentage of cells showing Arl13b in the cilia (C). (D and E) SEPTIN7 is localized in the cilia in Control KO cells while absent in SEPTIN9 KO cells expressing GFP-Centrin-DH-PH. (G and H) SEPTIN2 is localized in the cilia in Control KO cells and absent in SEPTIN9 KO cells expressing GFP-Centrin-DH-PH. (F and I) Percentage of cells showing SEPTIN2 and SEPTIN7 in the cilia in the Control KO cells and SEPTIN9 KO line transfected with GFP-Centrin-DH-PH. Data are represented as mean ± SEM. Student’s t-test p-value significance *=p<0.05, **=p<0.01, ***=p<0.001 and ****=p<0.0001. n.s. No significance is denoted. Bar represents 10 μm. Bars in magnified images represents 1 μm.

## Materials and Methods

### Cell Culture and Transfections

Immortalised human retinal pigment epithelial cells (hTERT-RPE1) were cultured in DMEM/F12 with 10% FBS and grown at 37°C with 5% CO2. Cells were serum-starved in DMEM/F12 for 16-24h to induce the formation of primary cilia. HeLa cells stably expressing the tetracycline-controlled transactivator protein (Tet-ON, Clontech) were cultured in DMEM with 10% FBS and grown at 37°C with 5% CO2. Cells were transfected with Lipofectamine 2000 (Invitrogen) using the manufacturer’s recommendations. Generally, 10µL of transfection reagent was used for every 4µg of plasmid per well of a 6-well tissue culture plate.

### Plasmid Construction and siRNA

For the binding studies and RhoA activation assays, all constructs are human-specific and use the CMV promoter. Generation of Myc-Arhgef18 is previously described. SEPTIN9_i3 is used as a representative SEPTIN9 isoform since it rescues cytokinesis defects in cells depleted of all SEPTIN9 isoforms (Estey et al., 2010). We designed a cloning system to enable easy shuttling of inserts between different vectors using two restriction enzymes: EcoRI and XhoI. The multiple cloning sites of Flag, Flag-His6 and GFP vectors were modified to contain EcoRI and XhoI sites that were in-frame with these affinity or epitope tags. The backbone for the Flag and Flag-His6 vectors is pcDNA3.0+ (Invitrogen), and that for the GFP vector is pEGFP-C1 (Takara Bio Inc).

Flag-His6-SEPTIN9, Flag-His6-SEPTIN9ΔN (268-568) and Flag-His6-RhoA were generated by subcloning or PCR amplifying inserts using EcoRI and XhoI overhangs. Flag-SEPTIN2, Flag-SEPTIN6, Flag-SEPTIN7 and Flag-SEPTIN9 were made by subcloning each septin using EcoRI and XhoI into a home-made Flag pcDNA3.0+ vector. GFP-SEPTIN9 was constructed by subcloning into the GFP vector using EcoRI and XhoI.

For lower protein expression, the CMV promoter from pEGFP-C1 (Takara Bio Inc) was removed and replaced with the SEPTIN2 promoter using AseI and AgeI. This vector was the basis for generating a low-expressing basal-body targeted construct. To target proteins to basal bodies, Centrin1 (Cetn1) was fused downstream of GFP. Cetn1 cDNA (generous gift from L. Pelletier, University of Toronto) was amplified by PCR using KpnI and EcoRI overhangs, and ligated into the low-expressing GFP vector. The resulting vector encoded GFP-Cetn1 protein expressed at lower amounts compared to using the CMV promoter. Low-expressing GFP-Cetn1-SEPTIN9 was made by subcloning SEPTIN9 using EcoRI and XhoI. GFP-Cetn1-DN (dominant negative) RhoA and GTP-Cetn1-DH-PH were generated by PCR amplification using EcoRI and XhoI overhangs. Mislocalised or severely over-expressed centrin-containing plasmids were excluded from analysis.

All siRNA sequences were obtained from Dharmacon. The siRNA sequence targeting human SEPTIN9 is 5’-GCACGAUAUUGAGGAGAAA. Individual Arhgef18 SmartPool siRNAs were screened for their ability to knockdown protein by immunofluorescence, and the most efficient siRNA was used for subsequent experiments including western blots. The siRNA sequence targeting human Arhgef18 is 5’-agaucaugcugaaggugua. All siRNAs were electroporated into hTERT-RPE1 using the Amaxa electroporator (Program X-01). All knockdown experiments used siRNA and electroporation, except for the quantitation for Fig. 3C and D. These experiments were transduced with lentiviral particles encoding an shRNA targeting human SEPTIN9 (V3LHS_317869, Dharmacon). In order to assess the functional implications of the interactions between the DH-PH domain of RhoGEF and RhoA on ciliogenesis in vivo, an argenine to alanine (R742A) mutation was generated in an N-terminal GFP and Centrin-tagged (GFP-Centrin-R742A DH-PH) construct using site directed mutagenesis (Strategene).

### Establishment of SEPTIN9-KO cell line using CRISPR/Cas9 system

The single-guide RNA (sgRNA) sequence targeting the human SEPTIN9 gene CCGGTGGACTTCGGCTACGTGGG was designed using CRISPOR (Zhang lab general cloning protocol) to knock out all SEPTIN9 isoforms (SEPTIN9_i1 through SEPTIN9_i5). Double-stranded oligonucleotides for the target sequence was inserted into the all-in-one sgRNA expression vector PX330-based plasmid, including SpCas9n D10A nickase+sgRNA. HeLa cells grown on a 12-well plate were transfected with the sgRNA vector using Lipofectamine 2000 Transfection Reagent (Thermo Fisher Scientific) as specified by the manufacturer. Western blot screening was used to identify putative knockout cells. The identified SEPTIN9 CRISPR Knockout (KO) cell line was subsequently analyzed using immunofluorescence and confocal microscopy.

### Protein Complex Pull-downs

HeLa Tet-ON (Clontech) cells were transfected using Lipofectamine 2000 (Invitrogen) on 6-well plates and harvested 18–24 h later. Cells were washed with cold PBS and lysed using buffer A (PBS, 20 mM imidazole, 1% Triton X-100, and a cOmplete® protease inhibitor tablet (Roche). Lysates were clarified for 30 min at 13,000 *g* at 4°C. The supernatant was then incubated with

Ni-NTA agarose (Qiagen) for 1 h and washed three times in buffer B (PBS, 20 mM imidazole, and 0.1% Triton X-100). SDS-PAGE loading buffer was added to beads, run on 10% SDS-PAGE gels, and immunoblotted with antibodies.

### RhoA Activation Assay

hTERT-RPE1 cells were transfected with Flag-His6-RhoA, Myc-Arhgef18 and/or the septin complex (Flag-SEPTIN2, Flag-SEPTIN6, Flag-SEPTIN7, GFP-SEPTIN9) on 6-well tissue-culture plates using Lipofectamine 3000 (Invitrogen) and harvested 18–24 h later. Cell lysates were harvested as outlined in the G-lisa RhoA Activation Assay Kit (Cytoskeleton Inc) using the supplied lysis buffer. Lysates were incubated in Rho-GTP affinity plate for 30 minutes. The plate contains a Rho-GTP-binding protein linked to the wells of the plate, which binds active GTP-bound Rho in cell lysates, while inactive GDP-bound Rho is removed during washing steps.

Subsequently, the bound active RhoA was detected with a RhoA specific antibody and chemiluminescence. Active and total RhoA levels were quantified on a Li-Cor Odyssey FC Imaging system.

### Western Blotting

Standard immunoblotting procedure was performed using 40 minute incubations of primary and secondary antibodies. Dilutions of antibodies used for immunoblotting were as follows: mouse anti-Flag at 1:7,500 (clone M2; Sigma-Aldrich), mouse anti-Myc at 1:1000 (9E10, Covance), rabbit anti-GFP at 1:20,000 (Invitrogen), mouse anti–glyceraldehyde 3-phosphate dehydrogenase at 1:50,000 (Millipore), rabbit anti-SEPTIN9 at 1:1,000(Surka et al., 2002) and rabbit anti-Arhgef18 (C-terminus) at 1:2000 (Nagata & Inagaki, 2005). HRP secondary antibodies were used at 1:5,000 (Jackson ImmunoResearch Laboratories, Inc).

### Immunofluorescence

For septin localisation along the length of cilia, cells were washed with room temperature PBS and then fixed in 1% PFA in PBS for 20 min at room temperature. PFA was inactivated and cells were permeabilized in buffer C (PBS with 25 mM glycine, 25 mM ammonium chloride, and 0.1% Triton X-100) for 20 min followed by blocking in PBS with 5% fetal calf serum for 20 min. Standard immunofluorescence procedures were used with 40 min incubations of primary and secondary antibodies with 3 washes in PBS+0.1% Tween-20. Dilutions of antibodies used for immunofluorescence were rabbit anti-SEPTIN9 at 1:100 (Surka et al., 2002), rabbit anti-Arhgef18 at 1:500 (Nagata & Inagaki, 2005), and mouse anti-acetylated tubulin at 1:1000 (6-11B-1, Sigma-Aldrich). Fluorescent secondary antibodies conjugated to AlexaFluor488 and AlexaFluor564 were used at 1:500 (Invitrogen). Coverslips were mounted on slides using fluorescence mounting medium (Dako).

For GFP-centrin localisation at the base of cilia, cells were briefly washed with room temperature PBS and then fixed with ice-cold methanol for 3 minutes. Cells were washed three times with PBS. The same blocking and immunostaining procedure as above was used. Rabbit anti-GFP (Invitrogen) and mouse anti-gamma tubulin (GTU-88; Sigma-Aldrich) was diluted at 1:1000 for immunofluorescence.

### Microscopy

Fixed cells were imaged at room temperature using an inverted fluorescence microscope (DMIRE2; Leica) equipped with an IEEE 1394–based digital camera (ORCA-ERG model C4742-95-12ERG; Hamamatsu Photonics). Images were acquired using a 63x or 100x oil immersion objective (HCX Plan Apochromat with numerical aperture of 1.40–0.7; Leica). The system was driven by Volocity acquisition software (PerkinElmer).

Live cells transfected with GFP-septins were imaged on a Quorum spinning disk confocal microscope. Coverslips were placed in a magnetic chamber and media was replaced with HPMI-buffered DMEM/F12 with 10% FBS. Live cells were imaged at room temperature using an inverted fluorescence microscope (DMIRE2, Leica) equipped with a Hamamatsu C9100-12 back-thinned EM-CCD camera and Yokogawa CSU 10 spinning disk confocal scan head (with Spectral Aurora Borealis upgrade). Two separate diode-pumped solid state laser lines were used at 491nm and 561nm (Spectral Applied Research) along with a 1.5x magnification lens. A 63x/1.4 objective was used with the following emission filters: 515 nm +/-40 and 594 nm +/-40. The system was operated with Perkin Elmer Volocity software.

### Statistical analysis

Two-tailed Student’s t tests and SuperPlots (Lord et al., 2020) were applied to determine statistical significance.

### Quantification of the fluorescence intensity of Exocyst and Transition zone proteins at the base of the cilia

The histogram graph in image J software was used to determine the threshold value of pixels in the entire image. In each image used in this study, all pixel values were in the range of 0-255 and no under or over saturated pixels were detected. In order to quantify the fluorescence intensity of the Exocyst proteins, Exo70 and Sec8, and the transition zone component, TCTN2 at the base of the cilia, a Region Of Interest (ROI) of 98.47 µm^3^, containing 24 pixels, was selected in all the fluorescence intensity measurements. The same ROI was also used to quantify the background fluorescence, subtracted from the Sum of ROI fluorescence intensity reads for each protein at the base of the cilia. Two-tailed Student’s t tests were applied to determine statistical significance.

## Notes

### Competing Interest Statement

The authors have declared no competing interest.

